# GMPR: A novel normalization method for microbiome sequencing data

**DOI:** 10.1101/112565

**Authors:** Li Chen, Jun Chen

## Abstract

**Summary:** Normalization is the first and a critical step in microbiome sequencing (microbiome-Seq) data analysis to account for variable library sizes. Though RNA-Seq based normalization methods have been adapted for microbiome-Seq data, they fail to consider the unique characteristics of microbiome-Seq data, which contain a vast number of zeros due to the physical absence or undersampling of the microbes. Normalization methods that specifically address the zeroinflation remain largely undeveloped. Here we propose GMPR - a simple but effective normalization method - for zeroinflated sequencing data such as microbiome-Seq data. Simulation studies and analyses of 38 real gut microbiome datasets from 16S rRNA gene amplicon sequencing demonstrated the superior performance of the proposed method.

**Availability and Implementation:** ‘GMPR’ is implemented in R andavailable at https://github.com/jchen1981/GMPR

**Supplementary Information:** Supplementary data are available at Bioinformatics online.

## 1 INTRODUCTION

Normalization, which aims to correct or reduce the bias introduced by variable library sizes (sequencing depths), is an essential step before any downstream statistical analyses for high-throughput sequencing experiments (e.g. RNA-Seq or microbiome-Seq) (Anders and Huber, 2010). An inappropriate normalization method may either introduce unwanted variation and hence reduce statistical power or, more severely, result in falsely discovered features. Normalization is especially critical when the library size is a confounding factor that correlates with the variable of interest. One popular approach for normalizing the sequencing data involves calculating a size factor for each sample as an estimate of the library size. The size factors can be used to divide the read counts to produce normalized data, or to be included as offsets in count-based regression models for differential feature analysis. One simple normalization method is TSS (Total Sum Scaling), which uses the total read count for each sample as the size factor. However, there are several undesirable properties for TSS. First, it is not robust to outlier counts. Outliers have frequently been observed in sequencing samples due to technical artifacts such as preferential amplification by PCR. Several outliers could bias the library size estimates significantly. Second, it creates compositional effects: non-differential features will appear to be differential due to the constant sum constraint. Compositional effects are much stronger for data where there are overly abundant features and the total number of features is relatively small. An ideal normalization method should thus capture the invariant part of the count distribution, and be robust to outliers and differential features. The latter property is important to reduce the false positives due to compositionality.

Many normalization methods have been developed primarily for RNA-Seq data. Popular RNA-Seq normalization methods include RLE (Relative Log Expression) implemented in the DESeq (Anders and Huber, 2010) and TMM (Trimmed Mean of M values) implemented in the edgeR (Robinson and Oshlack, 2010). Recently, the CSS (Cumulative Sum Scaling) method has been developed for microbiome Seq data (Paulson *et al.*, 2013). Compared to RNA-Seq data, microbiome-Seq data are more over-dispersed and contain a vast number of zeros. Take the COMBO data for example (Wu *et al.*, 2011), it contains 1,873 non-singleton OTUs (Operational Taxonomic Units, a proxy for bacterial species) from 98 samples with more than 90% of zeros. The observed zeros are a mixture of ‘structural zeros’ (due to physical absence) and ‘sampling zeros’ (due to undersampling). To circumvent the zeroinflation problem, one popular strategy is to add a pseudo-count. This practice has a Bayesian explanation and implicitly assumes that all the zeros are due to undersampling (McMurdie and Holmes, 2014). However, this assumption may not be appropriate due to the large extent of structural zeros. Therefore, a more rigorous method to address the zeroinflation in normalization is still much needed.

Here we propose a novel normalization method GMPR (Geometric Mean of Pairwise Ratios), developed specifically for zeroinflated sequencing data such as microbiome-Seq data. By comprehensive tests on simulated and real datasets, we show that GMPR outperforms the other competing methods for zeroinflated count data.

## 2 METHODS

Denote the *c_ki_* as the count of the *k*th OTU (*k* = 1, …, *q*)in the *i*th as (*i* = 1, …, *n*) microbiome-Seq sample. GMPR calculates the size factor *s_i_* for sample *i* as

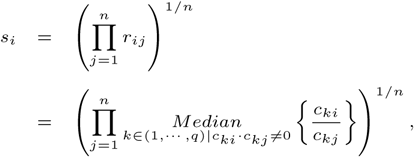

where *r_ij_* is the median count ratio on nonzero counts between sample *i* and *j*. Figure 1a illustrates the procedure of GMPR. The basic idea of GMPR is that we conduct the pairwise comparison first and then combine the pairwise results to obtain the final estimate. Using this strategy, we do not need to calculate the geometric mean for each OTU as implemented in RLE. Since only a small number of OTUs (or none) are shared across all samples due to severe zeroinflation, RLE is not stable or possible. However, for every pair of samples, they usually share many OTUs. Thus, for pairwise comparison, we focus on these common OTUs that are observed in both samples to have a reliable inference of the abundance ratio between samples. We then synthesize the pairwise abundance ratios using a geometric mean to obtain the size factor. To be noted, GMPR is a general method, which could be applied to any type of sequencing data in principle.

**Fig. 1.**
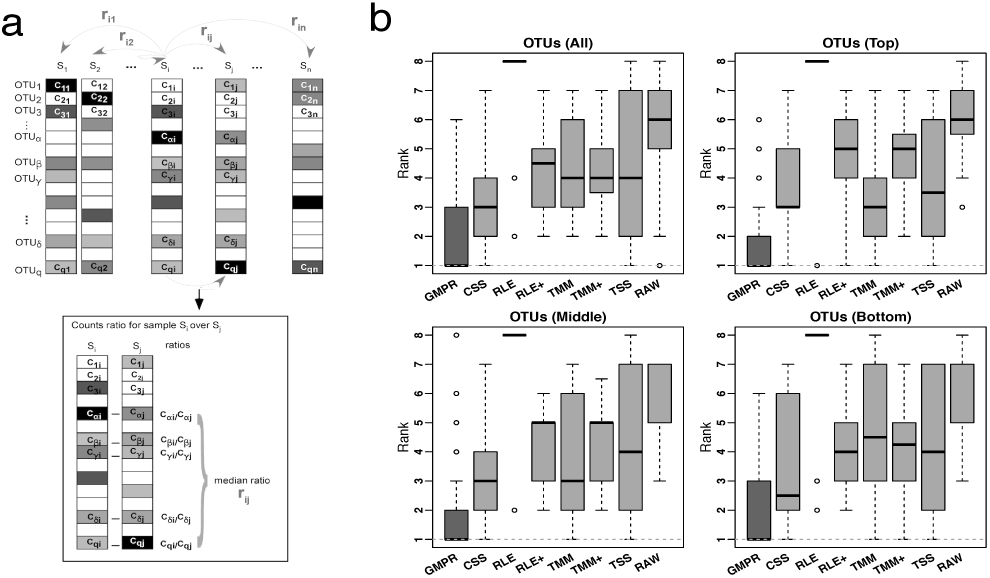
Illustration of the GMPR method and its performance on 38 real stool microbiome data sets. **a)** GMPR starts with pairwise comparisons (upper). Each pairwise comparison calculates the median abundance ratio on those common OTUs between the pair of samples (lower). The pairwise ratios are then synthesized into a final estimate. **b)** Distribution of the ranks for the medians of the variances over the 38 data sets. The median is calculated over all OTUs or OTUs of different prevalence level (Top, middle and bottom tertile)

## 3 RESULTS

We make use of both simulated and real microbiome-Seq data sets to compare GMPR to other normalization methods including CSS, RLE, RLE+ (RLE with pseudo-count 1), TMM, TMM+ (TMM with pseudo-count 1), and TSS. The details of how to estimate the size factors for each method could be found in Supplementary Note.

### 3.1 Simulation results

We use a perturbation-based simulation approach to evaluate the performance of normalization methods, focusing on their robustness to outlier OTUs and differentially abundant OTUs (Supplementary Figure S1). Briefly, we first generate zeroinflated count data based on a Dirichlet-multinomial model with known library sizes (Chen and Li, 2013). Next we ‘perturb’ the counts and evaluate the ability of a normalization method to recover the ‘true’ library sizes. We employ two perturbation approaches where we decrease/increase a‘fixed’ or ‘random’ set of OTUs with large fold changes. In the ‘fixed’ perturbation approach, the same set of OTUs are decreased/increased in the same direction for all samples, reflecting differentially abundant OTUs under a certain condition such as disease state. In the ‘random’ perturbation approach, each sample has a random set of OTUs perturbed with a random direction, mimicking the sample-specific outliers. Specifically, the mean and dispersion parameters of Dirichlet-multinomial distribution are estimated from the COMBO dataset after filtering out rare OTUs (prevalence < 10%) (Wu *et al.*, 2011). The library sizes are also drawn from those of the COMBO data. To investigate the effect of the amount of zeros, OTU counts are simulated with different zero percentages (~ 60%, 70% and 80%) by adjusting the dispersion parameter. A varying percentage of OTUs (0%, 1%, 2%, 4%, 8%, 16% and 32%) are perturbed in each set of simulation. We also study the effect of the strength of perturbation. The counts *c_ij_* of perturbed OTUs are changed to 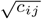 or 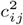 for strong perturbation, and 0.25*c_ij_* or 4*c_ij_* for moderate perturbation. Finally, size factors for all methods are estimated, and the Spearman's correlation between the size factors and ‘true’ library sizes is calculated. The simulation is repeated 50 times and the average Spearman's correlation is reported.

The simulation results are summarized in Supplementary Figure S2 and S3. It is not surprising that the performance of all methods trends to decrease with the increased zero percentage, number of perturbed OTUs and strength of perturbation. Interestingly, TSS has excellent performance under moderate perturbation but performs poorly under strong perturbation. GMPR, followed by CSS, consistently outperforms other methods when the perturbation is strong. When the perturbation is moderate, GMPR is only secondary to TSS when the percentage of zeros is high (80%), and on par with TSS when the percentage of zeros is moderate (70%) or low (60%). When the perturbed OTUs are random, the performance of RNA-Seq based methods such as TMM and RLE decreases significantly as the number of perturbed OTUs increases. Though the performance of RLE+ improves by adding pseudo-counts to RLE, the size factor estimated by TMM+ merely correlate with true library size when the zero percentage is moderate (70%) or high (80%). In contrast, GMPR, together with CSS, performs stable in all cases. Overall, we could see GMPR yields better size factor estimate than CSS.

### 3.2 Real datasets results

We next evaluate various normalization methods using 38 gut microbiome datasets from16S rRNA sequencing of the stool samples (Supplementary Table S1). These real datasets were retrieved from qiita database (https://qiita.ucsd.edu/) with a sample size larger than 50. The 38 datasets came from different species of both invertebrates and vertebrates as well as a wide range of biological conditions. We chose stool samples because the stool microbiota has been more studied than that from other body sites.

It is not feasible to calculate the correlation between estimated size factors and ‘true’ library sizes as done for simulations. Alternatively, we use the inter-sample variability as a performance measure since an appropriate normalization method will reduce the variability of the OTU counts due to different library sizes. A similar measure has been used in the evaluation of normalization performance for microarray data (Fortin *et al.*, 2014). We use traditional variance as the metric to assess inter-sample variability. For each method, the variance of each OTU across all samples is calculated, and the median of the variances of all OTUs or stratified OTUs (based on their prevalence) is reported for each study. For each study, all methods are ranked based on these median variances. The distributions of their ranks across these 38 studies for each method are depicted in Figure 1b. A higher ranking indicates that the method performs better in terms of minimizing inter-sample variability.

In Figure 1b, we could see that GMPR achieves the best performance with top ranks in 22 out of 38 datasets, followed by CSS, which tops in 7 datasets (Supplementary Table S2). This result is consistent with the simulation studies, where GMPR and CSS are overall more robust to perturbations than other methods. Moreover, GMPR performs consistently the best for reducing the variability of OTUs at different prevalence level. It is also noticeable that the inter-sample variability is the largest without normalization (RAW), and TSS does not perform well for a large number of studies. As expected, RLE only works for 8 out of 38 datasets due to a large percentage of zero read counts. By adding pseudo-counts, RLE+ improves the performance significantly compared to RLE. However, there is not much improvement of TMM+ compared to TMM. To see if the difference is significant, we performed paired Wilcoxon signed rank tests between the ranks of the 38 datasets obtained by GMPR and by any other methods. GMPR achieves significantly better ranking than other methods (p-value¡0.05 for all OTUs or stratified OTUs). Overall, GMPR achieves the best performance in terms of minimizing inter-sample variability.

